# Opposing seasonal dynamics in phylogenetic and flower color diversity of co-flowering wildflower assemblages

**DOI:** 10.1101/2025.10.21.683724

**Authors:** Christopher A. Talbot, Marjorie G. Weber

**Affiliations:** Department of Computational Biology, Cornell University, Ithaca, New York, USA; Department of Ecology & Evolutionary Biology, University of Michigan, Ann Arbor, Michigan, USA

**Keywords:** biodiversity, community science, co-occurrence, flower color, plant-pollinator interactions, community assembly, trait diversity, phenology

## Abstract

Flower color is a key trait mediating plant-pollinator interactions, yet the ecological rules governing its distribution across co-flowering assemblages remain poorly understood. We examined spatiotemporal patterns of flower color and phylogenetic diversity in Eastern North American wildflowers to test for non-random community structure, seasonal shifts, and associations between flower color similarity and phylogenetic relatedness. Using occurrence, phenology, and color data from community science datasets, we reconstructed communities of co-flowering assemblages and calculated standardized effect sizes of mean pairwise distances for both flower color and phylogenetic relatedness. We analyzed seasonal trends using mixed models and assessed whether closely related species co-flowered non-randomly with respect to color similarity using bootstrap correlations. We found that, on average, co-flowering assemblages were overdispersed for both flower color and phylogeny. However, flower color dispersion significantly decreased over the growing season, while phylogenetic dispersion significantly increased, reflecting a pattern in which co-flowering assemblages shifted to more distantly related but similarly colored assemblages. Finally, color and phylogenetic dispersion were negatively correlated across communities, indicating that communities composed of more closely related species tended to exhibit more dissimilar flowers. Together, our results suggest that wildflower assemblages in Eastern North America show detectable, non-random structure across space and time with respect to color and relatedness at the broad scale, and that mechanisms underlying the structure of co-flowering groups may shift throughout the growing season.

## 1 INTRODUCTION

Flowering plants display remarkable diversity in flower coloration both within and across co-flowering assemblages, a pattern that has long captivated biologists and naturalists (e.g., Lovell, 1909; Clegg & Durbin, 2000; Sobel & Streisfeld, 2013; Erickson & Pessoa, 2022). However, despite the large body of research on flower color evolution and ecology, the emergent patterns of color diversity across flowering plant communities remain understudied for most floras. Understanding large-scale patterns of flower color diversity can inform hypotheses about the emergent outcomes of mechanisms governing plant-pollinator interactions and community assembly processes. Here, we examine the spatiotemporal distribution of flower color diversity in common Eastern North American wildflowers, asking whether flower color shows non-random patterns of diversity at the community scale throughout the growing season.

Emergent patterns of flower color are hypothesized to be shaped by a suite of complex ecological and evolutionary factors. Most obviously, because flower color is a key phenotype in pollinator recognition and choice and the insurance of reproductive success in flowering plants, the flower color composition of plant communities should be mediated, at least in part, by pollinators. Pollinators can exhibit innate and learned color preferences (Ellis et al., 2021; Muñoz-Galicia et al., 2021) which could translate into predictable community-scale patterns in several ways. For example, if possessing distinct flower colors generally benefits wildflowers by attracting distinct pollinators and avoiding reproductive interference of hybridization from heterospecific pollen transfer (Muchhala et al., 2014), then an emergent pattern of flower color overdispersion within co-flowering assemblages may be expected. However, pollinator preference for flower color could also be hypothesized to drive flower color similarity, rather than dissimilarity. For example, in cases where pollinators are rare (de Jager et al., 2011), similar floral phenotypes may facilitate attracting higher pollinator densities, which could result in an emergent pattern of underdispersion in flower color. Pollinator diversity itself may also impact flower color diversity at the community scale: where or when pollinator diversity is high, there may be more flower color niches for species to occupy, facilitating flower color overdispersion (McEwen & Vamosi, 2010).

Another set of these hypotheses focuses on abiotic, rather than biotic, environmental conditions. In these hypotheses, the abiotic environment selects, filters, or plastically shifts species towards certain flower color phenotypes, leading to co-flowering assemblages clustered into fewer colors than expected by chance. Factors including temperature (Ahmad et al., 2022), UV irradiance (Koski & Ashman, 2016), and precipitation (Warren & Mackenzie, 2001) have been hypothesized to associate with flower color on regional scales. Because communities experience similar abiotic conditions, an overall trend of restricted flower color diversity at the community scale may be expected across communities (Patricia Willmer, 2011).

Phylogeny further complicates predictions of flower color diversity in communities. In flowering plants, closely related species tend to co-occur in space and time more frequently than expected by chance (“The Diversification of Flowering Plants through Time and Space”, 2006; Du et al., 2015; Simon et al., 2021). This pattern of phylogenetic relatedness may have implications for predictions of flower color similarity in co-flowering species. For example, the strength of several forces driving selection and filtering for divergent flower colors, such as competitive exclusion, reproductive interference, and deleterious hybridization, should be greater among close relatives than among distant ones. This interaction may lead to an emergent pattern in which communities composed of more closely related species have more dissimilar flower colors than expected by chance (i.e., phylogenetic dispersion would negatively correlate with flower color dispersion across communities). Previous studies have found flower color to be a labile trait with weak phylogenetic signal (Koski & Ashman, 2016; Tai et al., 2020), perhaps due to selection for color divergence between closely related, co-flowering species (Grossenbacher & Stanton, 2014; Milet-Pinheiro et al., 2021). However, studies examining specific plant clades and communities have reported both significant under-(e.g., Bergamo et al., 2020; de Jager et al., 2011) and over-dispersion (e.g., Makino & Yokoyama, 2015; McEwen & Vamosi, 2010; Skeels et al., 2021) in flower color.

Together, the suite of hypotheses about the drivers of flower color similarity generates conflicting predictions about emergent patterns across communities. It remains unclear whether an overall pattern will emerge across communities or whether the complex combination of drivers will result in a random color distribution at the broad scale, with no single mechanism emerging as a dominant driver. Further, patterns of flower color and phylogenetic structure may be highly context-dependent, varying across communities or throughout the growing season. To address these questions, we quantify the spatiotemporal patterns of flower color diversity across communities of common Eastern North American wildflowers and test whether predictions from ecological and evolutionary processes hold at the broad community scale. We ask (1) do co-flowering assemblages of wildflowers display non-random patterns of flower color across Eastern North America?

(2) Do patterns of flower color dispersion shift across the flowering season? And (3) how do changes in phylogenetic relatedness of co-flowering species relate to patterns of flower color dispersion over space and time? For the final question, we investigated patterns of relatedness at two scales: the whole-community scale and within closely related species pairs.

## 2 MATERIALS AND METHODS

All code for analysis and figure generation was written in R v4.4 and run in RStudio (Posit Team, 2025; R Core Team, 2024). Figure generation was performed using the ggplot2 package for R (Wickham et al., 2025).

A list of common Eastern North American flora was assembled using Newcomb’s Wildflower Guide (Newcomb, 1977). This field guide was selected for its thorough treatment of common wildflower species across Eastern North America. Taxonomy was standardized using Leipzig’s Catalogue of Vascular Plants and the lcvplants package (Freiberg et al., 2020). After standardization, subspecies were combined into a single parent species, and duplicates were removed.

For each species, occurrences within North America were retrieved from GBIF using the rgbif package for R (Chamberlain et al., 2024; GBIF.org, 2023a, 2023b, 2023c, 2023d, 2023e). Occurrences were filtered to include only those from later than 1960 and with less than 10 kilometers of uncertainty. We further filtered occurrences to ensure accuracy, removing those with geospatial issues (as detected by the GBIF API), coordinates in country or continent centroids, coordinates in oceans, and coordinates with a value of 0. All filtering steps were completed using custom download requests to the GBIF API. We created occurrence rasters comprising 10 km *×*10 km grid cells (100 km^2^ area) across North America using the terra package in R (Hijmans et al., 2025), which were then populated with presence/absence data for each species using the downloaded occurrences with speciesRaster package (now called epm) for R (Title et al., 2025).

To acquire phenology data, we downloaded iNaturalist occurrences annotated as in the “flowering” phenophase for each species from the Eastern North American region using the rinat package for R (Barve et al., 2022). We extracted the dates of each observation, removed data points *±>* 2 standard deviations from the median, and manually removed obvious outliers (Iwanycki Ahlstrand et al., 2022). Flowers were considered to be in bloom for the entire span from the earliest to the latest occurrence after cleaning.

We combined our presence/absence and phenology data to assemble groups of species that were flowering at the same time of year in the same geographic area (co-flowering assemblages). We randomly selected non-adjacent 10 km *×* 10 km grid cells (hereafter “sites”), creating a checker-board pattern of 1951 sites across the Northeast region. For each site, we extracted community lists of flowering species separately for each day from April 1 through November 1 (216 distinct days), creating a dataset of distinct co-flowering assemblages at each site for each day of the flowering season. In total, we acquired 216 days across our 1951 cells, yielding 421 267 co-flowering assemblages.

To acquire species color data, we created a custom program pyflocolor that uses a *k*-means clustering algorithm to identify *H, S*, and *V* color channel values from iNaturalist photographs. Images were downloaded from a random selection of 100 flowering occurrences for each species (or, all available images if there were fewer than 100 available). Using these images, we acquired the mean petal color of the most frequent distinctive color covering the greatest proportion of the petals using a modified version of a *k*-means clustering method from Perez-Udell et al. (2023). To train the algorithm, we manually selected the cluster containing flower petals for 10 initial images per species. Values of *k* were manually selected for each flower during this process. Variable *k* values were necessary due to the wide range of flower colors – in particular, flowers with colors more similar to their typical background required a higher *k* to differentiate. We then applied the HSV color range and *k* values from the manually selected clusters to obtain clusters for all other images of that species. The average HSV color of all flowers in the images was then computed and converted to the LAB color space. Color values were visually compared to representative flower images for each species to ensure accuracy. For color distance calculations, we calculated ΔE (perceptual color distance) between species pairs. Notably, our approach to quantifying color does not detect wavelengths in the ultraviolet part of the electromagnetic spectrum, which many pollinators can detect. Our analysis accounts for the visual color spectrum (roughly 400 to 700nm), which accounts for a large portion of color space and has been shown to be critically important for pollinator attraction (Spaethe et al., 2001).

We retrieved a phylogeny of our species by trimming tips from Smith & Brown’s vascular plant mega-tree (Smith & Brown, 2018). We calculated the phylogenetic distance of species pairs using the cophenetic() function in the ape package for R (Paradis et al., 2024). We calculated the phylogenetic signal of flower color using Blomberg’s *K* and Pagel’s *λ* statistics for each HSV and LAB color channel separately (Shrestha et al., 2014). These calculations were performed using the phylosig() function from the phytools package (Revell, 2025).

### 2.1 Patterns of phylogenetic and flower color dispersion across communities

To test whether co-flowering species in communities display non-random patterns in flower color and phylogenetic relatedness, we calculated the standardized effect size of mean pairwise distance (SES.MPD) for both flower color (HSV and LAB color space) and phylogenetic distances within each co-flowering assemblage using the picante package (Kembel et al., 2020). The SES.MPD metric quantifies the degree to which observed traits or phylogenetic distances deviate from a null model, with positive values indicating overdispersion and negative values indicating clustering.

**Figure 1.**
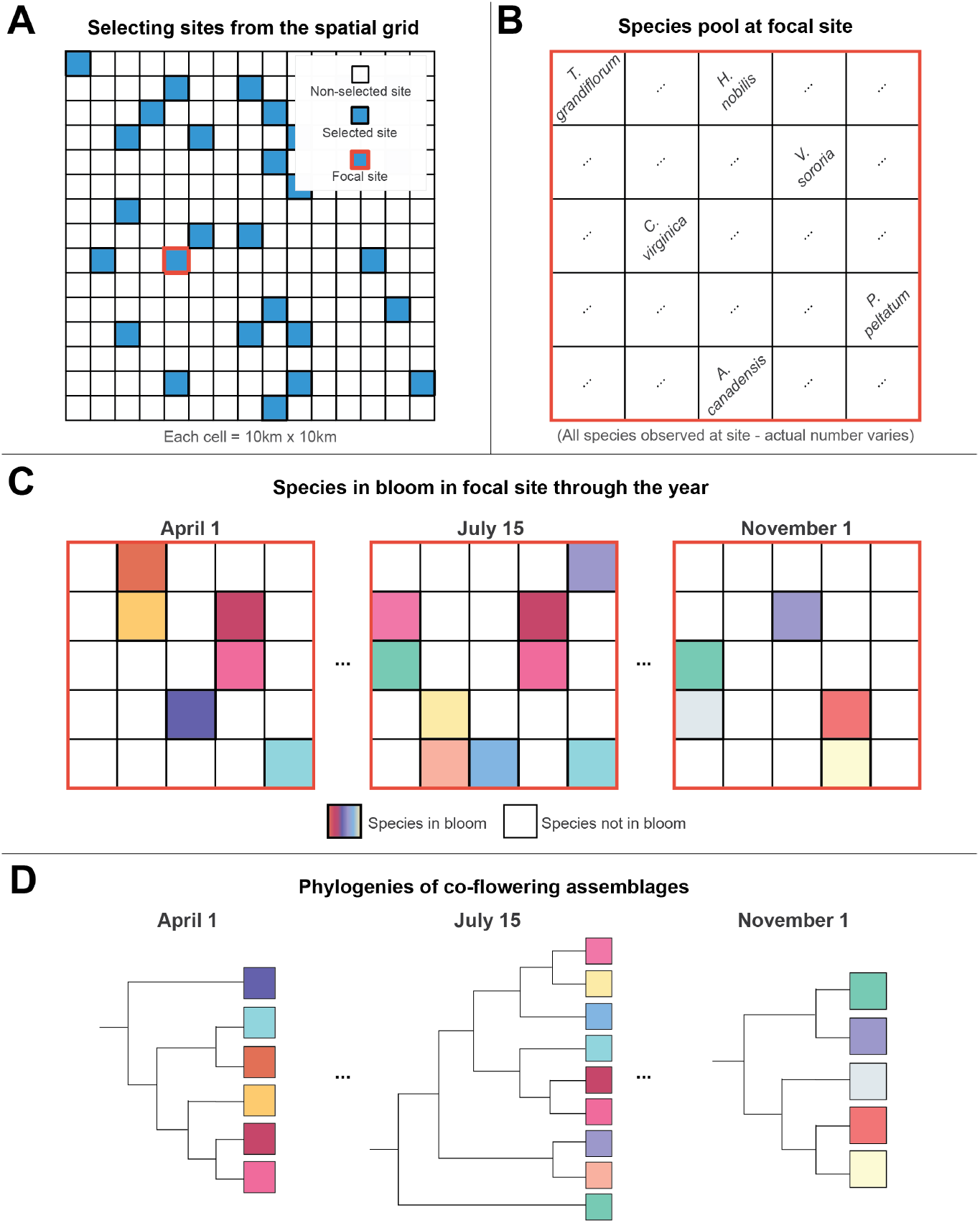
Schematic of sampling design for the study. We aggregated data from multiple sources to generate a large set of “co-flowering assemblages,” flowers blooming together in time and space, along with color and phylogeny data. **A**) Sites are 10 km*×*10 km grid cells across Eastern North America, selected randomly from the complete set of cells. The true number of sites selected was 1951. Here, we display a single focal site to demonstrate the data associated with a particular cell. **B**) At a given focal site, we compiled a list of all species observed using GBIF occurrence records. The number of species varies across sites. **C**) Given a set of species present at the focal site, we use phenology data from iNaturalist to generate “co-flowering assemblages,” the subset of flowers observed at the focal site and in bloom on a given day. We gathered color data for each species from iNaturalist using a custom pipeline. **D**) For each co-flowering assemblage, we calculated phylogenetic and flower color dispersion.

**Figure 2.**
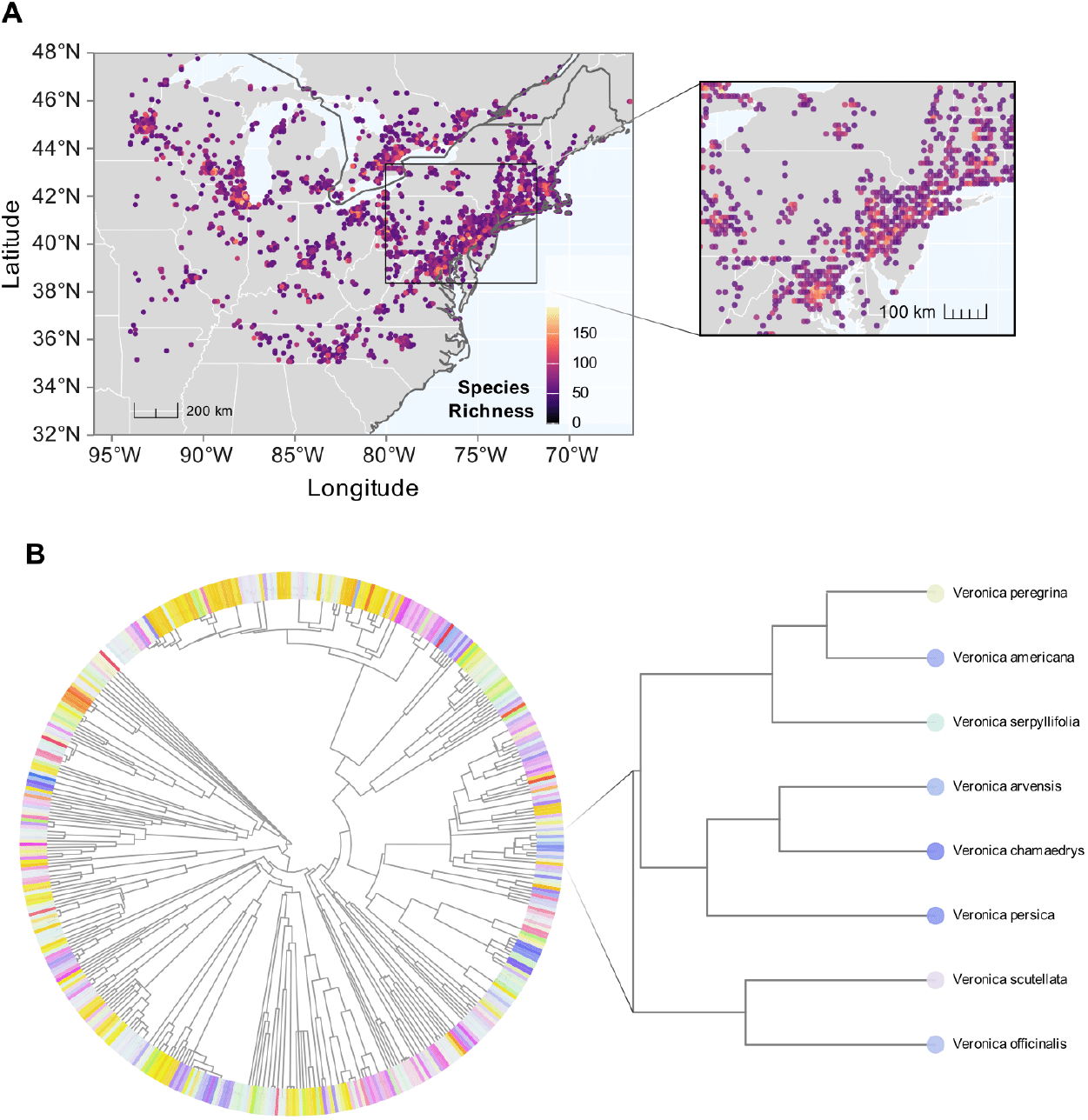
**A)** Research-grade species occurrence data were collected from GBIF. The Eastern North America region was divided into a raster of 10 km*×*10 km grid cells (100 km^2^ area); all grid cells in which that species was observed were identified. Cells were randomly selected, and cells bordering others were removed, resulting in 1951 cells in the final dataset. Here, we show the complete set of selected sites, colored by species richness within each site. Zooming in on a high-density region better displays the approximate cell size and checkerboard pattern (inset). **B)** The phylogeny of all species used in the study. Tip colors represent the assigned flower color for each species, identified using a *k*-means clustering algorithm from iNaturalist photographs. We zoom in on the *Veronica* genus with tips colored according to the calculated mean color for that species as an example of tip-level data.

For our null model, we used the ‘independentswap’ algorithm, which randomizes the community data matrix of each assemblage while preserving species richness per site and species occurrence frequency over the whole dataset. By preserving species richness and frequency while randomizing co-occurrence, we generate a null expectation for community assembly given the observed regional species pool of Eastern North America. This approach also accounts for the spatial and temporal structure of the data in the statistical test. We conducted 999 randomizations per community to generate robust null distributions. We then calculated the proportion of assemblages with significant overdispersion (SES.MPD*>*1.96) or clustering (SES.MPD*<−* 1.96) relative to the null distributions. To quantify the strength and direction of the overall pattern across co-flowering assemblages, we tested whether the distribution of SES.MPD values across assemblages significantly differed from zero using a one-sample t-test, and the mean SES.MPD was reported.

To directly test whether phylogenetic relatedness predicts flower color similarity across co-flowering assemblages while accounting for temporal and spatial structure, we fitted a generalized linear mixed model (GLMM) using the glmmTMB package (Brooks et al., 2025). The model included standardized flower color dispersion (z-transformed SES.MPD in the LAB color space) as the response variable, with standardized phylogenetic dispersion as the primary predictor. To account for temporal autocorrelation, we included time (day of year) as a covariate in the form of linear time and time squared as fixed effects. To account for the hierarchical and spatial autocorrelation of the data, we included random intercepts and random slopes for time nested within sites, as well as an exponential spatial correlation structure based on site coordinates that models the decay in similarity between sites with increasing geographic distance. We evaluated model fit using the Akaike Information Criterion (AIC) and calculated variance partitioning across site-level, spatial, and residual components to quantify the relative importance of geographic and temporal structure.

### 2.2 Seasonal patterns in composition

We next assessed whether patterns of flower color and phylogenetic dispersion shift throughout the growing season. To visualize patterns over time, we plotted the mean SES.MPD values for flower color and relatedness metrics across sites against the day of year. To test for temporal trends while accounting for the hierarchical structure of our data, we fitted linear mixed models for both phylogenetic and flower color dispersion using the `lme4` package (Bates et al., 2025). For each dispersion metric, we constructed four candidate models of increasing complexity: (1) a linear trend model using scaled day of year, (2) a quadratic trend model with an additional squared term, (3) a seasonal model using sine and cosine components to capture cyclical patterns, and (4) a combined model incorporating both linear trend and seasonal components. All models included random intercepts for sites to account for location-specific variation. We selected the most parsimonious model for each dispersion metric using the AIC. We calculated marginal R-squared values to assess the explanatory power of the fixed effects. To identify specific periods of clustering or overdispersion, we summarized patterns monthly by calculating the mean SES.MPD values with 95% confidence intervals for each month. The prevailing pattern (clustered, random, or overdispersed) for each month was assessed by determining whether the confidence intervals excluded zero.

### 2.3 Close relatives, co-occurrence, and flower color

Several key costs of co-flowering, such as hybridization, are hypothesized to apply to closely related species rather than to more distantly related pairs. As such, we performed analyses targeting only closely related species pairs. We defined closely related species pairs as those that diverged *≤*30 million years ago (mya), as this is approximately the mean time to the most recent common ancestor for modern flowering plant families (Magallón et al., 2015). Using only these species pairs, we conducted a bootstrap analysis with 1000 iterations that controlled for phylogenetic non-independence to ask whether species with similar flower colors co-flower less frequently than expected by chance. In each iteration, we randomly selected approximately 50 phylogenetically independent species pairs, selecting closely related pairs without replacement to ensure that each species occurred only once in a given iteration.

The spatial co-occurrence of each species pair was calculated using the Szymkiewicz-Simpson (overlap) coefficient, given the presence/absence raster (de Bello et al., 2016; Drury et al., 2018; Šizling et al., 2024). We calculated the color distance of each species pair using ^Δ^*E* in the LAB color space. For each bootstrap iteration, we calculated the Spearman rank correlation between spatial overlap and color distance. To analyze patterns across all bootstrap iterations, we calculated the mean correlation coefficient, the proportion of iterations with statistically significant correlations, and the proportion showing negative correlations (suggesting that more similar colors correspond to less spatial overlap). We also divided species pairs into quartiles based on color similarity to examine whether the relationship between color similarity and spatial overlap followed a linear or non-linear pattern.

To evaluate whether closely related species pairs display patterns distinct from distantly related pairs, we repeated our bootstrap analysis using distantly related species pairs – selecting approximately 50 species pairs *≥* 250 mya diverged – to compare with recently diverged species pairs.

## 3 RESULTS

Flower color, quantified in the LAB color space, exhibited a weak but statistically significant phylogenetic signal. Blomberg’s *K* values were 0.011 (*P* = 0.007) for *L* (lightness), 0.008 (*P* = 0.025) for the *a*-channel (green-red), and 0.025 (*P* = 0.001) for the *b*-channel (blue-yellow). Similarly, Pagel’s *λ* values were 0.434 (*P <* 0.001), 0.650 (*P <* 0.001), and 0.809 (*P <* 0.001) for *L, a*, and *b* channels, respectively, indicating that closely related species tend to be slightly less similar in flower color than expected under a null Brownian Motion model of trait evolution.

### 3.1 Patterns of flower color and phylogenetic dispersion

Co-flowering assemblages, on average, showed statistically significant deviation from random expectations for both flower color and phylogenetic dispersion. The mean standardized effect size of the mean pairwise distance (SES.MPD) for flower color (LAB) was slightly positive (mean SES.MPD = 0.010, 95% CI [0.009, 0.010]; *t*_420 005_ = 30.35, *P <* 0.001), suggesting a tendency towards flower color overdispersion across all communities. Similarly, phylogenetic relatedness within assemblages showed a slight, statistically significant tendency towards overdispersion (mean SES.MPD = 0.006, 95% CI [0.006, 0.007]; *t*_420 005_ = 18.62, *P <* 0.001). Despite these significant overall means, the vast majority of individual co-flowering assemblages (99.7% for both flower color and phylogenetic relatedness) exhibited patterns of dispersion that were not significantly different from random expectations (i.e., SES.MPD values between *−*1.96 and 1.96). Our spatial-temporal generalized linear mixed model revealed a significant negative relationship between phylogenetic and flower color dispersion across co-flowering assemblages (*β* =*−* 0.079, SE = 0.002, *z* =*−*50.02, *P <* 0.001). Variance partitioning revealed substantial geographic structure in patterns of flower color dispersion. Site-level variation accounted for 36.3% of total variance, spatial autocorrelation explained 26.7%, and residual variation comprised 37.0%.

### 3.2 Seasonal trends in community assembly

Both flower color and phylogenetic dispersion exhibited significant seasonal variation. Linear mixed models with site as a random effect best explained these temporal patterns when incorporating both linear and cyclical (sine and cosine of the day of year) terms.

For flower color dispersion (LAB), the best-fitting model (selected by AIC; marginal *R*^2^ = 0.001, conditional *R*^2^ = 0.356) revealed a significant negative linear trend with season progression (scaled day of year: estimate = *−*0.018, *P <* 0.001), indicating that flower colors within assemblages became, on average, more similar (less dispersed) as the growing season advanced. Significant cyclical components were also detected (sin(day): estimate = *−*0.017, *P <* 0.001; cos(day): estimate = 0.011, *P <* 0.001).

Consistent with these results, our spatial-temporal generalized linear mixed model confirmed a significant quadratic effect of time on color dispersion (*β* = 0.027, SE = 0.001, *z* = 27.10, *P <* 0.001). Random slopes for time varied substantially across sites (SD =0.505), indicating considerable heterogeneity in seasonal patterns across the landscape.

Conversely, phylogenetic dispersion showed a significant positive linear trend. The best-fitting model (AIC; marginal *R*^2^ = 0.007, conditional *R*^2^ = 0.391) indicated that co-flowering species became, on average, more distantly related (more phylogenetically overdispersed) as the growing season progressed (scaled day of year: estimate = 0.040, *P <* 0.001). A significant sine component also contributed to this seasonal pattern (sin(day): estimate = 0.030, *P <* 0.001). In contrast, the cosine component was not significant (cos(day): estimate = 0.001, *P* = 0.108). The observed monthly mean SES.MPD values showed considerable variation, often with confidence intervals overlapping zero, highlighting that the subtle seasonal trends detected by the mixed models emerged when accounting for site-specific effects and analyzing the continuous progression of the season.

### 3.3 Close relatives, co-occurrence, and flower color similarity

We investigated whether flower color similarity predicted co-occurrence across closely related species pairs (diverged *≤*30 mya). A bootstrap analysis of phylogenetically independent species pairs (1000 iterations, average 50 pairs per iteration) yielded a mean Spearman rank correlation (*ρ*) between spatial overlap and LAB color distance of *−*0.045 (SD = 0.091), reflecting a trend for species pairs with similar colors to display less spatial overlap. However, while 69.1% of bootstrap iterations resulted in negative correlations, only 4.8% of these correlations were statistically significant (*P <* 0.05), indicating that this trend was not statistically significant.

We found that this trend was further weakened in distantly related species pairs (diverged *≥*250 mya), where the bootstrap analysis of phylogenetically independent species pairs (1000 iterations,average 50 pairs per iteration) yielded a mean Spearman rank correlation (*ρ*) between spatial overlap and LAB color distance of *−*0.018 (SD = 0.097). For these distantly related pairs, only 56.7% of bootstrap iterations resulted in a negative correlation, and only 3% of the correlations were statistically significant (*P <* 0.05).

**Figure 3.**
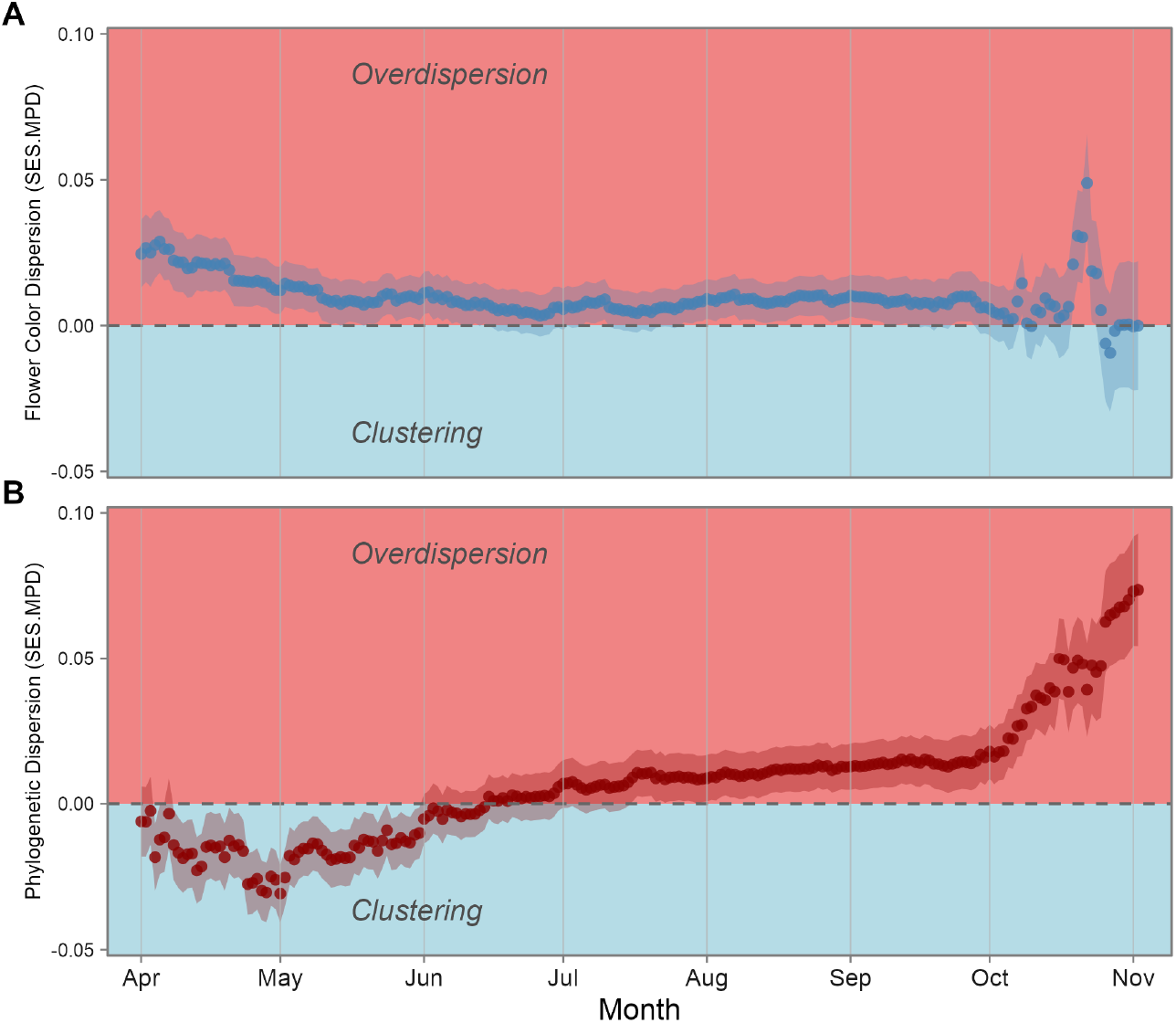
Seasonal trends in mean phylogenetic and flower color dispersion over the growing season. Individual points represent the mean observed flower color or phylogenetic dispersion across all co-flowering assemblages on a particular day of the year. The highlighted region represents the 95% confidence interval around the mean. **A**) Flower color dispersion shows a negative trend over time, transitioning from a more overdispersed state (with greater color variety) to a more random or clustered state (with less color variety). **B**) Phylogenetic dispersion displays an opposing trend when compared to flower color dispersion. Co-flowering assemblages display more phylogenetic clustering (closer relatives) early in the growing season, transitioning to more phylogenetic overdispersion (more distant relatives) later in the growing season.

When examining spatial overlap across quartiles of flower color similarity (using one representative bootstrap sample of 50 pairs of close relatives), species pairs with the most similar flower colors (*Q*_1_) exhibited the lowest mean spatial overlap (mean = 0.058, SE = 0.015). In contrast, pairs with moderately different colors (*Q*_2_) showed higher mean spatial overlap (mean = 0.116, SE = 0.022). However, an ANOVA comparing mean spatial overlap across the four color similarity quartiles did not find a statistically significant overall difference (*F*_3,96_ = 1.29, *P* = 0.282). A post-hoc Tukey HSD test also showed no significant difference in spatial overlap between *Q*_1_ and *Q*_2_ (difference = 0.058, 95% CI [*−*0.019, 0.135], *P*_adj_ = 0.210).

**Figure 4.**
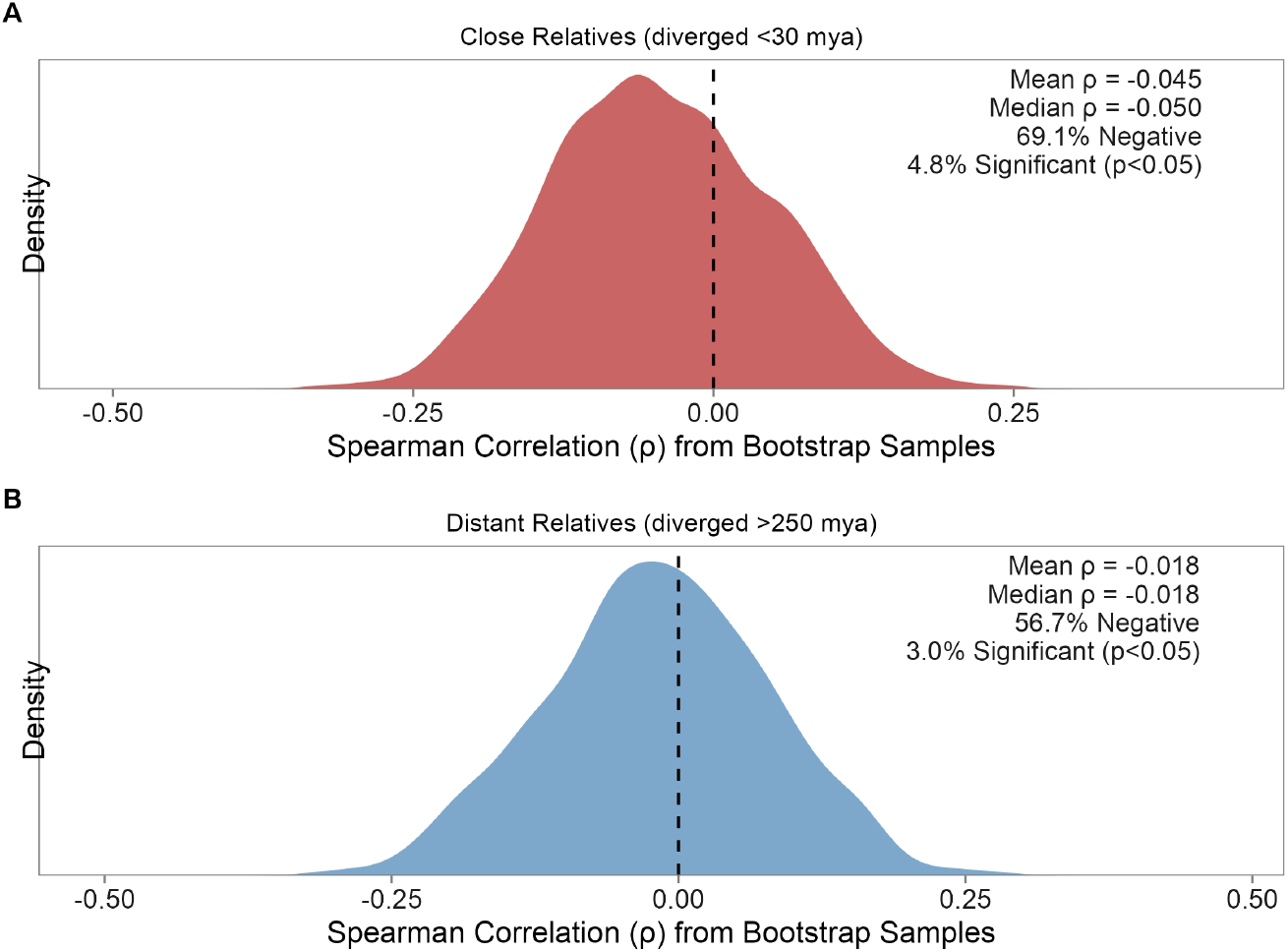
**A**) The density of Spearman correlation calculations across all bootstrap samples for phylogenetically independent, closely related species pairs (diverged *≤*30 mya). The mean relationship between flower color distance and spatial overlap is skewed towards negative values (indicating that greater similarity in flower color tends to result in less spatial overlap), but the distribution overlaps strongly with zero. **B**) The density of Spearman correlation calculations across all bootstrap samples for phylogenetically independent, distantly related species pairs (diverged *>* 250mya). In contrast to close relatives, distant relatives show a weaker tendency toward a negative relationship.

## 4 DISCUSSION

The diversity of colors found in co-flowering plant communities is hypothesized to be shaped by a complex interplay of ecological and evolutionary factors, making it unclear whether patterns should emerge at broad spatial scales. Here, we analyzed co-flowering plant assemblages across Eastern North American wildflower communities using data from large-scale community science projects. Our study revealed three main findings. First, on average, co-flowering assemblages were significantly overdispersed for both flower color and phylogenetic relatedness across the region. Second, the two axes of diversity we examined exhibited opposing seasonal trajectories: flower color became more clustered as the season progressed, while phylogenetic relatedness became more overdispersed. And third, communities composed of more closely related species exhibited more dissimilar flower colors. We discuss each of these findings in light of the broader literature, along with their caveats and future directions.

### 4.1 Co-flowering Assemblages were Overall Overdispersed for Color and Relatedness

Across the entire dataset, we detected a signal of overdispersion for both flower color and phylogeny. Flower color overdispersion is generally expected to be the outcome of pollinator-mediated competition, where selection should favor divergent colors to reduce reproductive interference and hybridization and enhance pollinator constancy (Grossenbacher & Stanton, 2014; Muchhala et al., 2014). Similarly, phylogenetic overdispersion is often interpreted as a signature of limiting similarity, where competition for shared resources excludes close relatives due to niche overlap (Cavender-Bares et al., 2004; Kraft et al., 2007). Previous studies have found mixed evidence for both flower color and phylogenetic dispersion across co-flowering species. For example, McEwen and Vamosi (2010) found that co-flowering species in five subalpine Canadian meadows were overdispersed for color but not relatedness. Similarly, LeCroy et al. (2021) found that across serpentine seep communities in California, co-flowering assemblages were overdispersed for color but showed no significant phylogenetic structure. Yet, Shrestha et al. (2019) found no significant trend towards flower color or phylogenetic overdispersion or clustering.

Notably, while we found a statistically significant overall pattern of overdispersion across our dataset, individual assemblages ranged from significantly overdispersed to significantly underdispersed, with many indistinguishable from random. On the one hand, the emergence of an overall pattern of overdispersion at the regional scale, despite variation across individual assemblages, highlights the power of broad macroecological perspectives to detect signals that can be lost at smaller scales. On the other hand, this variation suggests that the forces shaping the emergent patterns of diversity are likely context-dependent or diluted by the stochastic nature of colonization and extinction across a large and heterogeneous geographic area (Menéndez-Serra et al., 2023; Viana & Chase, 2019). Other work has found that variation in color dispersion across communities can be predictably tied to factors such as community size (LeCroy et al., 2021), or shifting abiotic and biotic pressures across environmental gradients (Shrestha et al., 2014). Future work in this system, focused on quantifying these factors through climatic and field pollinator surveys, would be a natural next step to the large-scale pattern-based work presented here.

### 4.2 Opposing Seasonal Dynamics

The most compelling and unexpected finding of our study was the opposing trends in flower color and phylogenetic structure observed across the growing season. As the season progressed, co-flowering assemblages became increasingly phylogenetically overdispersed while simultaneously becoming more clustered in terms of flower colors. Together, this suggests that the interplay of factors that shape flower color diversity in communities changes throughout the growing season.

The trend toward increasing phylogenetic overdispersion directly challenges the prevailing view that late-season co-flowering assemblages are structured primarily by strong environmental filters that favor species from clades adapted to more challenging conditions (Ndiribe et al., 2013; Veldhuisen et al., 2025). For instance, recent work in European grasslands by Veldhuisen et al. (2025) found that late-season communities became more phylogenetically clustered, likely as abiotic pressures favored the success of tolerant lineages like *Asteraceae*. Our results for Eastern North America suggest a different primary driver. One plausible explanation is that early spring communities are phylogenetically constrained, dominated by specialist clades adapted to cold soils and short growing windows (e.g., spring ephemerals in clades such as *Liliaceae* and *Ranunculaceae*). As the season progresses and a wider array of environmental niches becomes available, species from more disparate lineages can establish and flower, leading to a net increase in phylogenetic dispersion before the ultimate filter of winter terminates the growing season in the region.

Simultaneously, we observed a decrease in flower color dispersion, indicating that communities became more homogeneous in color later in the season. Two non-exclusive mechanisms could drive this shift towards color clustering. First, abiotic filtering or selection may favor certain pigments that offer physiological benefits under late-season conditions, such as higher UV radiation or changing temperatures (Ahmad et al., 2022; Koski & Ashman, 2016). Second, as pollinator abundance or diversity declines late in the season (Bishop et al., 2024), plants may experience selection for convergence onto the most common or attractive floral signal, a form of community-wide Müllerian mimicry or facilitation that enhances collective attraction (de Jager et al., 2011; Johnson et al., 2003). The convergence on common colors, such as yellow and white, in late-season composites (e.g., *Solidago, Symphyotrichum*) across North America may be a visible manifestation of this phenomenon.

### 4.3 Phylogenetic Relatedness Negatively Correlates with Flower Color Dispersion Across Communities

Our spatial-temporal mixed model analysis provides direct evidence that phylogenetic and flower color dispersion are significantly and inversely related across co-flowering assemblages of Eastern North American wildflower species. Our analysis reveals a pattern where co-flowering assemblages composed of more closely related species displayed greater flower color dissimilarity across communities, while accounting for spatial and seasonal variation.

Together, a negative correlation between color dispersion and phylogenetic diversity means that sites with higher color diversity cannot be explained simply by the addition of more distantly related species. Several specific mechanisms could underlie this broader pattern. One potential explanation could be that closely related species in a community have experienced stronger selection for color divergence to avoid deleterious hybridization and reproductive interference in sympatry (Grossenbacher & Stanton, 2014; Muchhala et al., 2014). Such character displacement in sym-patric, closely related species has been documented in clades such as *Phlox* (Hopkins & Rausher, 2012) and *Mimulus* (Schemske & Bradshaw, 1999). The evolutionary lability of flower color may facilitate divergence among closely related species and convergence among distantly related co-flowering lineages (van der Niet & Johnson, 2012). Indeed, our results add to a substantial body of evidence pointing to flower color as an evolutionarily labile trait that can evolve rapidly among closely related species pairs (Koski & Ashman, 2016; Tai et al., 2020).

Further, the substantial spatial structure we detected, with over 60% of variance attributable to site identity and spatial autocorrelation, suggests that local and regional processes play a considerable role in shaping flower color diversity. Geographic variation in pollinator communities, climate, or biogeographic history may create regional tendencies toward specific phenotypes, with neighboring sites showing greater similarity due to shared environmental conditions. Future work linking the macroevolutionary patterns of flower color at the clade scale with macroecological patterns of diversity will help untangle the relative contributions of divergence among close relatives, potential convergence among distantly related species, and spatially structured drivers of community composition.

Despite the region-wide negative correlation between flower color dispersion and phylogenetic diversity, we did not find strong evidence that similarly colored species co-flowered more or less frequently than expected by chance at the pairwise scale. Our bootstrap analysis of closely related (*≤*30 mya) species pairs revealed only a weak, non-significant trend for similarly colored species to co-occur less frequently than expected. Consistent with expectations, this signal was even weaker for distantly related pairs (*≥* 250 mya), for whom the risks of hybridization and reproductive interference are reduced, but neither signal was significant. This pattern is consistent with other literature showing that flower color dissimilarity does not predict range overlap in closely related plant species (e.g., Weber et al., 2018), and suggests that our region-wide patterns were not driven by closely related species pairs with similar colors sorting into different communities.

### 4.4 Limitations and Future Directions

Our study leverages the power of community science data, allowing us to test for broad-scale patterns across a large phylogenetic and geographic range. However, this approach also has inherent limitations. For one, our approach for quantifying color from photographs captures only the human visual color spectrum. Importantly, this omits color data in the UV spectrum that is important for pollinator signaling, including patterning and nectar guides, and data cannot be interpreted through pollinator visual models (Chittka & Menzel, 1992). Further, community science photographs are unstandardized, which introduces noise into the data. As such, studies using flower color data from photographs are inherently limited, and - like any work using an incomplete floral trait set - hypotheses regarding pollinator perceptions should be interpreted in light of a complex suite of floral signaling traits (e.g., UV spectrum color, patterning, flower shape, nectar, and volatiles). However, despite these limitations, quantifying the color similarity of flowers from photographs in the visual spectrum remains valuable. As it reflects a substantial component of ecologically important flower color variation (Spaethe et al., 2001), it allows for large sample sizes in analyses of broad-scale patterns. It is of general interest because it is the component of flower color variation perceptible to the human eye. Regardless, pairing large-scale studies of photographs with finer-scale work that enables full signal analysis of flowers is critical for determining the drivers of color diversity across sites.

Other limitations of the scale of our study are that species presences are based on observations, rather than systematic surveys, and our definition of “co-flowering” assumes flowering throughout a species’ observed temporal range, which may not capture finer-scale phenological variations over space and time. Further, our approach does not capture within-species variation in color. However, despite these limitations, the detection of large-scale patterns highlights the value of these datasets for macroecological inquiry (Cavender-Bares et al., 2004; Kraft et al., 2007; Menéndez-Serra et al., 2023; Viana & Chase, 2019).

This work lays the foundation for more targeted research. Future studies should integrate local pollinator community data to directly test whether shifts in pollinator guilds drive the observed seasonal trends in flower color. Experimental manipulations that alter the color composition of artificial communities could directly test the mechanisms of facilitation versus competition. Finally, exploring these patterns in other biogeographic regions with different evolutionary histories and across environmental gradients will be crucial to determining the generality of the seasonal dynamics we have uncovered in Eastern North America.

## ACKNOWLEDGMENTS

The authors thank Nathan Sanders and James Boyko for helpful insight on methods and analyses; Matthew Hack for guidance on spatial and phylogenetic methods; Stephen Smith for advice on defining closely and distantly related species pairs; the Weber Lab for reviews of manuscripts and discussions on methods; and the University of Michigan Undergraduate Research Opportunity Program for support and funding throughout portions of this study.

## AUTHOR CONTRIBUTIONS

**Christopher A. Talbot**: Conceptualization (equal); data curation (lead); formal analysis (lead); investigation (lead); methodology (equal); visualization (lead); writing – original draft (equal); writing – review & editing (equal); funding acquisition (supporting). **Marjorie G. Weber**: Conceptualization (equal); investigation (supporting); methodology (equal); supervision (lead); visualization (supporting); writing – original draft (equal); writing – review & editing (equal); funding acquisition (lead).

## References

Ahmad, S., Chen, J., Chen, G., Huang, J., Zhou, Y., Zhao, K., Lan, S., Liu, Z., & Peng, D. (2022). Why Black Flowers? An Extreme Environment and Molecular Perspective of Black Color Accumulation in the Ornamental and Food Crops. Frontiers in Plant Science, 13. 10.3389/fpls.2022.885176

Barve, V., Hart, E., & Guillou, S. (2022, June). Rinat: Access ‘iNaturalist’ Data Through APIs.

Bates, D., Maechler, M., Bolker [aut, B., cre, Walker, S., Christensen, R. H. B., Singmann, H., Dai, B., Scheipl, F., Grothendieck, G., Green, P., Fox, J., Bauer, A., copyright on simulate.formula), P.N.K. (, Tanaka, E., Jagan, M., & Boylan, R. D. (2025, March). Lme4: Linear Mixed-Effects Models using ‘Eigen’ and S4.

Bergamo, P. J., Streher, N. S., Wolowski, M., & Sazima, M. (2020). Pollinator-mediated facilitation is associated with floral abundance, trait similarity and enhanced community-level fitness. Journal of Ecology, 108(4), 1334–1346. 10.1111/1365-2745.13348

Bishop, G. A., Fijen, T. P. M., Raemakers, I., van Kats, R. J. M., & Kleijn, D. (2024). Bees go up, flowers go down: Increased resource limitation from late spring to summer in agricultural landscapes. Journal of Applied Ecology, 61(3), 431–441. 10.1111/13652664.14576

Brooks, M., Bolker, B., Kristensen, K., Maechler, M., Magnusson, A., McGillycuddy, M., Skaug, H., Nielsen, A., Berg, C., van Bentham, K., Sadat, N., Lüdecke, D., Lenth, R., O’Brien, J., Geyer, C. J., Jagan, M., Wiernik, B., Stouffer, D. B., Agronah, M., … Krieger, N. (2025, October). glmmTMB: Generalized Linear Mixed Models using Template Model Builder.

Cavender-Bares, J., Ackerly, D. D., Baum, D. A., & Bazzaz, F. A. (2004). Phylogenetic Overdispersion in Floridian Oak Communities. The American Naturalist, 163(6), 823–843. 10.1086/386375

Chamberlain, S., Oldoni, D., Barve, V., Desmet, P., Geffert, L., Mcglinn, D., Ram, K., rOpenSci (https://ropensci.org/), Waller [aut, J., & cre. (2024, September). Rgbif: Interface to the Global Biodiversity Information Facility API.

Chittka, L., & Menzel, R. (1992). The evolutionary adaptation of flower colours and the insect pollinators’ colour vision. Journal of Comparative Physiology A, 171(2), 171–181. 10.1007/BF00188925

Clegg, M. T., & Durbin, M. L. (2000). Flower color variation: A model for the experimental study of evolution. Proceedings of the National Academy of Sciences of the United States of America, 97(13), 7016–7023. 10.1073/pnas.97.13.7016

de Bello, F., Fibich, P., Zelený, D., Kopecký, M., Mudrák, O., Chytrý, M., Pyšek, P., Wild, J., Michalcová, D., Sádlo, J., Šmilauer, P., Lepš, J., & Pärtel, M. (2016). Measuring size and composition of species pools: A comparison of dark diversity estimates. Ecology and Evolution, 6(12), 4088–4101. 10.1002/ece3.2169

de Jager, M. L., Dreyer, L. L., & Ellis, A. G. (2011). Do pollinators influence the assembly of flower colours within plant communities? Oecologia, 166(2), 543–553. 10.1007/s00442-010-1879-7

The Diversification of Flowering Plants through Time and Space: Key Innovations, Climate and Chance. (2006). In T.R. Hodkinson & J. A. N. Parnell (Eds.), Reconstructing the Tree of Life. CRC Press.

Drury, J. P., Tobias, J. A., Burns, K. J., Mason, N. A., Shultz, A. J., & Morlon, H. (2018). Contrasting impacts of competition on ecological and social trait evolution in songbirds. PLoS Biology, 16(1), e2003563. 10.1371/journal.pbio.2003563

Du, Y., Mao, L., Queenborough, S. A., Freckleton, R. P., Chen, B., & Ma, K. (2015). Phylogenetic constraints and trait correlates of flowering phenology in the angiosperm flora of China. Global Ecology and Biogeography, 24(8), 928–938. 10.1111/geb.12303

Ellis, A. G., Anderson, B., & Kemp, J. E. (2021). Geographic Mosaics of Fly Pollinators With Divergent Color Preferences Drive Landscape-Scale Structuring of Flower Color in Daisy Communities. Frontiers in Plant Science, 12, 617761. 10.3389/fpls.2021.617761

Erickson, M. F., & Pessoa, D. M. A. (2022). Determining factors of flower coloration. Acta Botanica Brasilica, 36, e2021abb0299. 10.1590/0102-33062021abb0299

Freiberg, M., Winter, M., Gentile, A., Zizka, A., Muellner-Riehl, A. N., Weigelt, A., & Wirth, C. (2020). LCVP, The Leipzig catalogue of vascular plants, a new taxonomic reference list for all known vascular plants. Scientific Data, 7(1), 416. 10.1038/s41597-020-00702-z

GBIF.org. (2023a, November). GBIF Occurrence Download. 10.15468/dl.vs5hv2

GBIF.org. (2023b, November). GBIF Occurrence Download. 10.15468/dl.sn77ag

GBIF.org. (2023c, November). GBIF Occurrence Download. 10.15468/dl.ghv5fm

GBIF.org. (2023d, November). GBIF Occurrence Download. 10.15468/dl.fpt9wv

GBIF.org. (2023e, November). GBIF Occurrence Download. 10.15468/dl.cfafx8

Grossenbacher, D. L., & Stanton, M. L. (2014). Pollinator-mediated competition influences selection for flower-color displacement in sympatric monkeyflowers. American Journal of Botany, 101(11), 1915–1924. 10.3732/ajb.1400204

Hijmans, R. J., Barbosa, M., Bivand, R., Brown, A., Chirico, M., Cordano, E., Dyba, K., Pebesma, E., Rowlingson, B., & Sumner, M. D. (2025, May). Terra: Spatial Data Analysis.

Hopkins, R., & Rausher, M. D. (2012). Pollinator-Mediated Selection on Flower Color Allele Drives Reinforcement. Science, 335(6072), 1090–1092. 10.1126/science.1215198

Iwanycki Ahlstrand, N., Primack, R. B., & Tøttrup, A. P. (2022). A comparison of herbarium and citizen science phenology datasets for detecting response of flowering time to climate change in Denmark. International Journal of Biometeorology, 66(5), 849–862. 10.07/s00484-022-02238-w

Johnson, S. D., Peter, C. I., Nilsson, L. A., & Ågren, J. (2003). Pollination Success in a Deceptive Orchid Is Enhanced by Co-Occurring Rewarding Magnet Plants. Ecology, 84(11), 2919– 2927. 10.1890/02-0471

Kembel, S. W., Ackerly, D. D., Blomberg, S. P., Cornwell, W. K., Cowan, P. D., Helmus, M. R., Morlon, H., & Webb, C. O. (2020, June). Picante: Integrating Phylogenies and Ecology.

Koski, M. H., & Ashman, T.-L. (2016). Macroevolutionary patterns of ultraviolet floral pigmentation explained by geography and associated bioclimatic factors. New Phytologist, 211(2), 708–718. 10.1111/nph.13921

Kraft, N. J. B., Cornwell, W. K., Webb, C. O., & Ackerly, D. D. (2007). Trait Evolution, Community Assembly, and the Phylogenetic Structure of Ecological Communities. The American Naturalist, 170(2), 271–283. 10.1086/519400

LeCroy, K. A., Arceo-Gómez, G., Koski, M. H., Morehouse, N. I., & Ashman, T.-L. (2021). Floral Color Properties of Serpentine Seep Assemblages Depend on Community Size and Species Richness. Frontiers in Plant Science, 11. 10.3389/fpls.2020.602951

Lovell, J. H. (1909). The Color Sense of the Honey-Bee: Is Conspicuousness an Advantage to Flowers? The American Naturalist, 43(510), 338–349. 10.1086/279064

Magallón, S., Gómez-Acevedo, S., Sánchez-Reyes, L. L., & Hernández-Hernández, T. (2015). A metacalibrated time-tree documents the early rise of flowering plant phylogenetic diversity. New Phytologist, 207(2), 437–453. 10.1111/nph.13264

Makino, T. T., & Yokoyama, J. (2015). Nonrandom Composition of Flower Colors in a Plant Community: Mutually Different Co-Flowering Natives and Disturbance by Aliens. PLoS ONE, 10(12), e0143443. 10.1371/journal.pone.0143443

McEwen, J. R., & Vamosi, J. C. (2010). Floral colour versus phylogeny in structuring subalpine flowering communities. Proceedings of the Royal Society B: Biological Sciences, 277(1696), 2957–2965. 10.1098/rspb.2010.0501

Menéndez-Serra, M., Ontiveros, V. J., Cáliz, J., Alonso, D., & Casamayor, E. O. (2023). Understanding stochastic and deterministic assembly processes in microbial communities along temporal, spatial and environmental scales. Molecular Ecology, 32(7), 1629–1638. 10.1111/mec.16842

Milet-Pinheiro, P., Santos, P. S. C., Prieto-Benítez, S., Ayasse, M., & Dötterl, S. (2021). Differential Evolutionary History in Visual and Olfactory Floral Cues of the Bee-Pollinated Genus Campanula (Campanulaceae). Plants, 10(7), 1356. 10.3390/plants10071356

Muchhala, N., Johnsen, S., & Smith, S. D. (2014). Competition For Hummingbird Pollination Shapes Flower Color Variation in Andean Solanaceae. Evolution, 68(8), 2275–2286. 10.1111/evo.12441

Muñoz-Galicia, D., Castillo-Guevara, C., & Lara, C. (2021). Innate and learnt color preferences in the common green-eyed white butterfly (Leptophobia aripa): Experimental evidence. PeerJ, 9, e12567. 10.7717/peerj.12567

Ndiribe, C., Pellissier, L., Antonelli, S., Dubuis, A., Pottier, J., Vittoz, P., Guisan, A., & Salamin, N. (2013). Phylogenetic plant community structure along elevation is lineage specific. Ecology and Evolution, 3(15), 4925–4939. 10.1002/ece3.868

Newcomb, L. (1977). Newcomb’s Wildflower guide (1st ed.). Little, Brown. Open Library ID: OL4534415M.

Paradis, E., Blomberg, S., Bolker [aut, B., cph, Brown, J., Claramunt, S., Claude, J., Cuong, H. S., Desper, R., Didier, G., Durand, B., Dutheil, J., Ewing, R. J., Gascuel, O., Guillerme, T., Heibl, C., Ives, A., Jones, B., Krah [aut, F., … de Vienne, D. (2024, December). Ape: Analyses of Phylogenetics and Evolution.

Patricia Willmer. (2011). Pollination and Floral Ecology. Princeton University Press.

Perez-Udell, R. A., Udell, A. T., & Chang, S.-M. (2023). An automated pipeline for supervised classification of petal color from citizen science photographs. Applications in Plant Sciences, 11(1), e11505. 10.1002/aps3.11505

Posit Team. (2025). RStudio: Integrated Development Environment for R.

R Core Team. (2024). R: A language and environment for statistical computing.

Revell, L. J. (2025, January). Phytools: Phylogenetic Tools for Comparative Biology (and Other Things).

Schemske, D. W., & Bradshaw, H. D. (1999). Pollinator preference and the evolution of floral traits in monkeyflowers (Mimulus). Proceedings of the National Academy of Sciences, 96(21), 11910–11915. 10.1073/pnas.96.21.11910

Shrestha, M., Dyer, A. G., Bhattarai, P., & Burd, M. (2014). Flower colour and phylogeny along an altitudinal gradient in the Himalayas of Nepal. Journal of Ecology, 102(1), 126–135. 10.1111/1365-2745.12185

Shrestha, M., Dyer, A. G., Garcia, J. E., & Burd, M. (2019). Floral colour structure in two Australian herbaceous communities: It depends on who is looking. Annals of Botany, 124(2), 221–232. 10.1093/aob/mcz043

Simon, A. D. F., Marx, H. E., & Starzomski, B. M. (2021). Phylogenetic restriction of plant invasion in drought-stressed environments: Implications for insect-pollinated plant communities in water-limited ecosystems. Ecology and Evolution, 11(15), 10042–10053. 10.1002/ece3.7776

Šizling, A. L., Keil, P., Tjørve, E., Tjørve, K. M. C., Žárský, J. D., & Storch, D. (2024, December). Mathematically and biologically consistent framework for presence-absence pairwise indices. 10.1101/2021.07.14.452244

Skeels, A., Dinnage, R., Medina, I., & Cardillo, M. (2021). Ecological interactions shape the evolution of flower color in communities across a temperate biodiversity hotspot. Evolution Letters, 5(3), 277–289. 10.1002/evl3.225

Smith, S. A., & Brown, J. W. (2018). Constructing a broadly inclusive seed plant phylogeny. American Journal of Botany, 105(3), 302–314. 10.1002/ajb2.1019

Sobel, J. M., & Streisfeld, M. A. (2013). Flower color as a model system for studies of plant evodevo. Frontiers in Plant Science, 4, 321. 10.3389/fpls.2013.00321

Spaethe, J., Schmidt, A., Hickelsberger, A., & Chittka, L. (2001). Adaptation, constraint, and chance in the evolution of flower color and pollinator color vision. Cambridge University Press. Accepted: 2011-06-01T14:28:42Z.

Tai, K.-C., Shrestha, M., Dyer, A. G., Yang, E.-C., & Wang, C.-N. (2020). Floral Color Diversity: How Are Signals Shaped by Elevational Gradient on the Tropical–Subtropical Mountainous Island of Taiwan? Frontiers in Plant Science, 11, 582784. 10.3389/fpls.2020.582784

Title, P., Swiderski, D., & Zelditch, M. (2025, February). Epm: EcoPhyloMapper.

van der Niet, T., & Johnson, S. D. (2012). Phylogenetic evidence for pollinator-driven diversification of angiosperms. Trends in Ecology & Evolution, 27(6), 353–361. 10.1016/j.tree.2012.02.002

Veldhuisen, L. N., Enquist, B. J., & Dlugosch, K. M. (2025, March). Phylogenetic diversity of flowering plants declines across the growing season in Rocky Mountain wildflower communities. 10.1101/2023.11.06.565878

Viana, D. S., & Chase, J. M. (2019). Spatial scale modulates the inference of metacommunity assembly processes. Ecology, 100(2), e02576. 10.1002/ecy.2576

Warren, J., & Mackenzie, S. (2001). Why are all colour combinations not equally represented as flower-colour polymorphisms? New Phytologist, 151(1), 237–241. 10.1046/j.1469-8137.2001.00159.x

Weber, M. G., Cacho, N. I., Phan, M. J. Q., Disbrow, C., Ramírez, S. R., & Strauss, S. Y. (2018). The evolution of floral signals in relation to range overlap in a clade of California Jewelflowers (Streptanthus s.l.) Evolution; International Journal of Organic Evolution, 72(4), 798–807. 10.1111/evo.13456

Wickham, H., Chang, W., Henry, L., Pedersen, T. L., Takahashi, K., Wilke, C., Woo, K., Yutani, H., Dunnington, D., van den Brand, T., Posit, & PBC. (2025, April). Ggplot2: Create Elegant Data Visualisations Using the Grammar of Graphics.

